# ERICA: Emulated Retinal Image CApture - A tool for testing, training and validating retinal image processing methods

**DOI:** 10.1101/2021.02.22.432253

**Authors:** Laura K Young, Hannah E Smithson

## Abstract

High resolution retinal imaging systems, such as adaptive optics scanning laser ophthalmoscopes (AOSLO), are increasingly being used for clinical and fundamental studies in neuroscience. These systems offer unprecedented spatial and temporal resolution of retinal structures *in vivo*. However, a major challenge is the development of robust and automated methods for processing and analysing these images. We present ERICA (Emulated Retinal Image CApture), a simulation tool that generates realistic synthetic images of the human cone mosaic, mimicking images that would be captured by an AOSLO, with specified image quality and with corresponding ground truth data. The simulation includes a self-organising mosaic of photoreceptors, the eye movements an observer might make during image capture, and data capture through a real system incorporating diffraction, residual optical aberrations and noise. The retinal photoreceptor mosaics generated by ERICA have a similar packing geometry to human retina, as determined by expert labelling of AOSLO images of real eyes. In the current implementation ERICA outputs convincingly realistic *en face* images of the cone photoreceptor mosaic but extensions to other imaging modalities and structures are also discussed. These images and associated ground-truth data can be used to develop, test and validate image processing and analysis algorithms or to train and validate machine learning approaches. The use of synthetic images has the advantage that neither access to an imaging system, nor to human participants is necessary for development.

## Introduction

Cellular-resolution imaging of the human retina *in vivo* has revolutionised retinal research, providing access to structures and functional processes in the retina, and it is increasingly being used in clinical research^1–6^. By enabling finer spatial and temporal scale measurements, adaptive optics (AO) retinal imaging offers the potential for closer monitoring of disease^7,8^. Furthermore, as an optically accessible piece of neural tissue that also shares its vascular origin with the cerebral vascular system, the retina is an important target for studying both neurological^9^ and vascular disorders^10^.

However, as with many biomedical imaging techniques, one challenge is the bottleneck imposed by the processing steps that are required to generate high quality images and extract information from them. Given the cellular-resolution that can be achieved with the use of adaptive optics, any image processing and analysis methods should operate with similar or better precision to fully utilise this capability. Furthermore, it is important to consider how the accuracy of the processing and analysis methods that are used impacts the output measures.

The approach taken here is to use synthetic data generated from a physically plausible model of image capture, which offers a number of advantages. Firstly, the properties of the simulated data capture can be systematically manipulated to investigate the effects of different image characteristics (e.g. quality) on the outcome measures, allowing us to understand the sources of error and to define tolerance limits. The parameters of the simulation (e.g. the field of view, frame rate and pixel size) can also be adjusted to simulate different instruments and investigate differences between systems. Secondly, a major advantage of using synthetic images is that they can be generated (and labeled) more rapidly and reliably than can human-derived images. Simulation therefore allows the production of the very large data sets that are typically required for machine learning approaches. Finally, synthetic data sets allow image processing and analysis methods to be developed and tested without needing access to a physical system, to human participants, or to experienced manual image graders. This more easily opens up the image processing challenge to the computer vision community. We first summarise the technological advances in high resolution retinal imaging, then we introduce the data capture and image processing pipeline, including current bottlenecks, and specific use-cases.

## Background

Imaging has long been an essential tool for studying the retina and techniques such as fundus imaging, scanning laser ophthalmoscopy and optical coherence tomography play a crucial role in the clinical diagnosis and management of ophthalmic disease. Miller *et al* 1996^11^ showed that cellular resolution imaging is possible outside the foveal center in some young eyes with good optical quality, provided that low order aberrations (defocus and astigmatism) are corrected. However, the application of AO techniques to retinal imaging, first attempted by Liang *et al*. in 1997^12^, led to a step change in imaging resolution by additionally compensating for the inherent higher-order optical aberrations in the eye, beyond defocus and astigmatism.

AO retinal imaging provides unprecedented opportunities for fundamental vision research^5,13^. Cellular resolution imaging allows the study of basic physiological processes in the retina. When combined with functional methods, such as single-cell stimulation^14^, blood flow measurements^15^ or recording intrinsic responses of individual photoreceptors^16,17^, the neurophysiology of the visual system can be studied in fine detail. AO has enabled precise mapping of the trichromatic cone mosaic^18^ and study of the neural wiring from individual photoreceptors to downstream neurons in the visual system^19–21^. When combined with psychophysical approaches, structural and functional AO retinal imaging provides new capabilities for studying visual perception on a cellular scale. For example, retinal imaging (with and without AO correction) has been used for measuring the microscopic eye movements made during fixation and understanding the role that they play in spatial vision^22–27^. This is due to the superior spatial and temporal resolution offered by AO imaging and because these systems measure the movement of the stimulus over the photoreceptors directly. In particular, AO retinal imaging offers tracking and stabilised stimulation with sub-cellular accuracy^28^. This is typically an order of magnitude better than a dual Purkinje eye tracker^29–31^ and two orders of magnitude better than a high resolution video-based eye tracker^32^. AO-enabled vision research not only advances our fundamental understanding of the healthy visual system, but it also provides a route to developing valuable cellular biomarkers of disease.

At present the most commonly used AO retinal imaging system is the AOSLO^5^, while AO flood illumination is used by many groups and AO optical coherence tomography systems are rapidly being developed and used. Pioneered by Roorda *et al*. in 2002^33^, the AOSLO generates images by raster-scanning a point source over an area of the retina and measuring the reflected light after focusing it through a confocal pinhole. This provides high lateral and axial resolution, but eye movements during the scan introduce motion distortions in the images. These motion distortions must be estimated and removed so that individual frames can be registered and averaged to produce an image with a high signal to noise ratio. A useful outcome of this process is the cellular-precision measurement of eye motion that is obtained. Motion estimation relies on selection of a motion-free reference frame against which to compare individual raw frames. The extraction algorithm needs to be robust to changes in image quality and to large and fast movements that can can cause significant distortion. Errors at this stage will reduce the quality of the resulting image and they will propagate through later image processing and analysis steps. After motion extraction, registration and averaging, single images obtained at adjacent retinal locations are often stitched together to form a larger field of view montage. Descriptive statistics, such as cone density estimates or reflectance changes, are often extracted from single or montage images. Image pre-processing steps, such as filtering, may also be used at any stage.

### AOSLO data analysis

There is a drive in the field to widen the adoption of AO retinal imaging instruments^5^, but there are challenges that are still to be overcome. As an example, Cooper *et al*.^34^ showed that motion distortions in AOSLO images can affect the reliability and accuracy of cone mosaic metrics derived from those images, and this could have clinically-relevant implications. The image processing and data extraction steps for AOSLO images often require some manual intervention, such as manual reference frame selection or the manual labeling of cone cells. Automated and semi-automated approaches for reference frame selection^35^, motion extraction and registration^36–39^ and montaging^40,41^ have been described. Many automated and semi-automated methods^36,40,42–46^ as well as machine learning approaches^47,48^ for cone localisation have been, and continue to be, explored. Assessing the accuracy of image processing and data analysis methods requires ground truth data, which is difficult or often impossible to obtain from real images captured through the imaging system. Another difficulty for machine learning approaches is that large data sets are typically required for training and validation. Even with efficiency improvements, AOSLO data capture with human participants and the manual generation of ground truth data is time consuming.

Accuracy estimation using ground truth data derived from real images is possible. Motion estimation accuracy has previously been assessed by imaging a model eye and introducing known or independently-measured movements^49^. Methods of cone identification have also been validated against semi-automated or manually labeled images^44^, though this is limited by the accuracy of the labels. A useful alternative approach is to use synthetic images for which ground truth data are specified. Such an approach has been demonstrated for testing the accuracy of motion extraction from an AOSLO using synthetic data created by resampling real images^50,51^. Synthetic images have also been used to test the accuracy of cone detection algorithms^52,53^ and, recently, to train deep learning models for image enhancement^54^. These approaches have generated synthetic retinal mosaics by randomly placing cells^52^ or by modifying a hexagonal packing matrix^53,55,56^ but stop short of simulating the full data capture pipeline.

### ERICA

We present ERICA (Emulated Retinal Image CApture), the first end-to-end simulation of AOSLO data capture generating a realistic output, with specified ground truth data, comparable to the stream of images obtained from human participants. ERICA has three main stages to generate fully synthetic images with plausible spatial and temporal structure:

1. A self-organising retinal cone mosaic generated with ground truth cone locations, reflectances, sizes and intensity profiles.
2. A simulation of eye movements added to the raster scan to create a spatio-temporal sampling pattern.
3. Image generation that replicates data capture through an AOSLO, including diffraction, noise and residual wavefront error.

Image processing and analysis algorithms are commonly applied to images of the photoreceptor mosaic, though approaches applied to split detection imaging have also been described^41,57,58^. ERICA generates images of the cone mosaic, but we also suggest extensions to the simulation to include rods in the mosaic, to add blood vessels and to emulate non-confocal imaging. It would also feasible to extend the model to 3D imaging, based on cellular-resolution retinal mapping in depth and an estimate of the 3D point spread function (PSF) of the eye and optical system.

## Methods and technical detail

ERICA is written in the Python programming language^59,60^ and operates in three stages, outlined in detail in the following sections and summarised in Figure 1. Firstly, a retinal mosaic is generated using a self-organising approach and this defines the location of cone cells (and thus provides the ground truth for cone localisation). The retinal mosaic is then defined parametrically (location, size, shape and reflectance of each cone) and an image of the ground truth retina is generated based on these parameters. The second component of the model is a simulation of the natural eye movements made during fixation. These movements are superimposed on the motion of the raster scanning system to define a spatio-temporal sampling pattern. Thirdly, for each temporal sample, the data capture through an AOSLO is modeled, including diffraction, optical aberrations and noise. Repeating this data capture simulation for each retinal position sampled produces a temporal sequence, analogous to the signals from a photodetector in an AOSLO, which can be processed as a synthetic frame. These procedures are repeated for each frame and the output is a set of images (which can be converted to a video stream, if desired) that approximate those captured from a human participant. Here we outline the current implementation of the three main modules, but alternative approaches and extensions to the simulation can be implemented within the ERICA framework.

**Figure 1.**
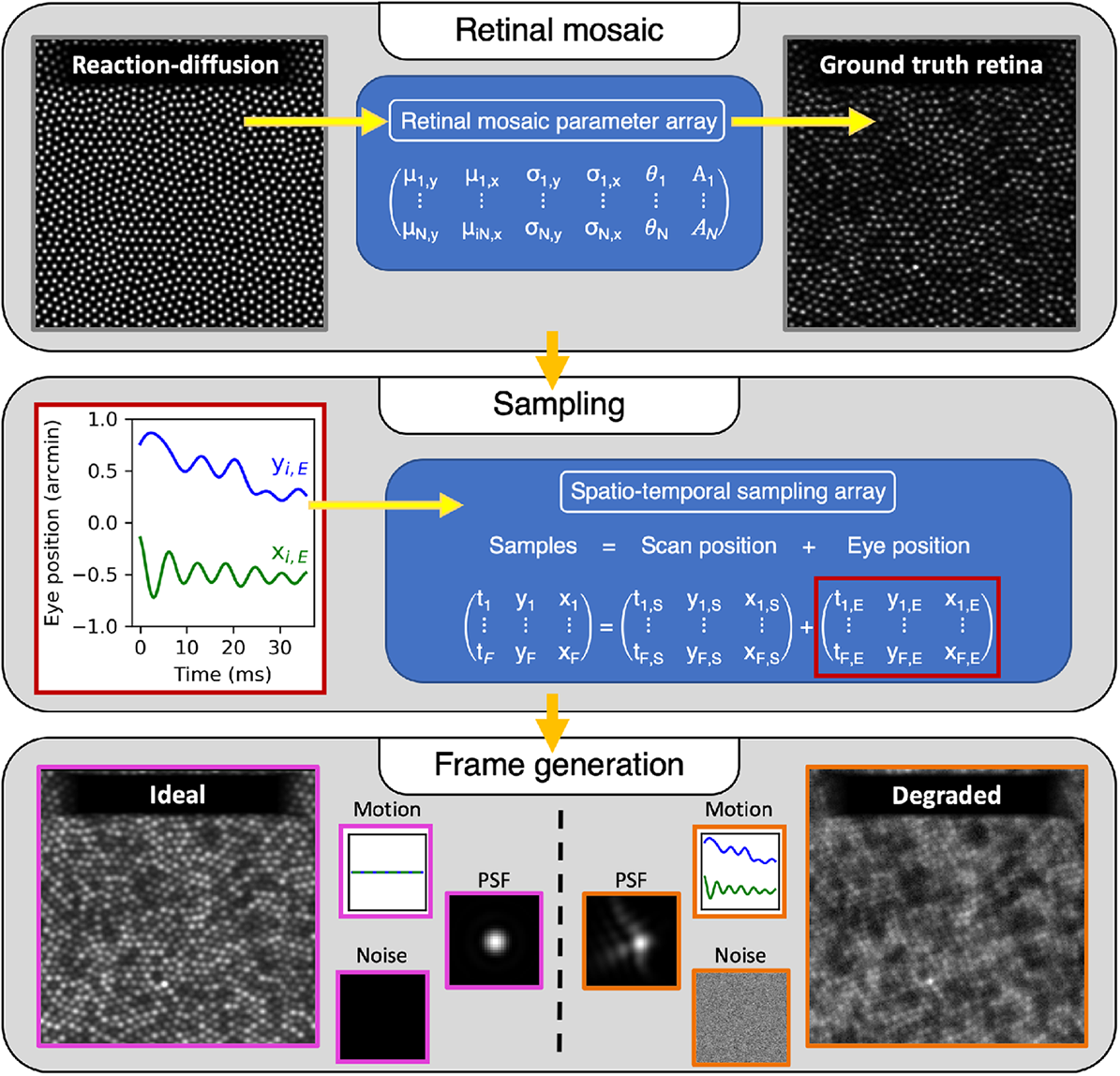
A schematic representation of the three main stages of ERICA. First, a retinal mosaic (left) is generated using reaction-diffusion modelling and the cone positions are extracted. Each of the N cones is defined parametrically as an elliptical Gaussian blob with parameters *µ*_*y*_ (vertical coordinate), *µ*_*x*_ (horizontal coordinate), *σ*_*y*_ (vertical width), *σ*_*x*_ (horizontal width), *A* (reflectance), and *θ* (orientation), and these are saved in the retinal mosaic parameter array. Second, eye position data are generated and added to the scan position data to define how the retina will be sampled. Third, frames are generated, as described in Figure 4, incorporating eye motion, diffraction and residual optical aberrations (the PSF) and noise. The example on the left is a diffraction limited image with no noise or eye motion and can be used as a ground truth frame. The example on the right includes plausible eye motion (drift and tremor, in this case), a realistic PSF based on aberration measurements^61^, and noise based on measurements through a real system.

### Self-organising retinal mosaic

The first stage of the simulation is to generate a realistic mosaic of cone cells. While such a synthetic mosaic has been generated by other means^52,53,55,56,62^, our approach is to use a self-organising method based on reaction-diffusion modelling to define the layout of the cone cells.

#### Cone locations

Reaction-diffusion systems model a process by which two chemicals (U and V) each diffuse and react with one another. Such systems have been used to generate complex self-organising patterns, also known as Turing patterns^63^, and have been offered as an explanation for the intricate pigmentation patterns formed in the coats of animals, such as leopards. The Gray-Scott reaction-diffusion model is one approach for generating these self-organising patterns and corresponds to the following irreversible chemical reactions that produce an inert product *P*:

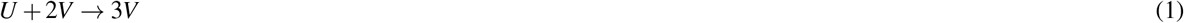

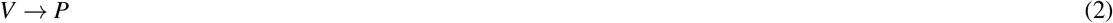

The model computes the concentrations, *u* and *v*, of the two chemicals *U* and *V* over a series of time steps using the following reaction-diffusion equations^64^:

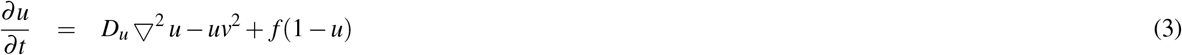

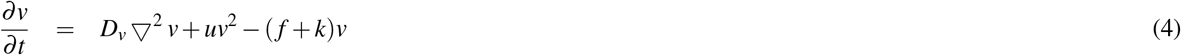

where *D*_*v*_ is the diffusion coefficient for chemical V, *D*_*u*_ is the diffusion coefficient for chemical U, *f* is the rate at which chemical U is replenished (Equation 1) and *k* is the rate at which the excess chemical V is removed (Equation 2). To produce a pattern of spots (visible as the final concentration of V) the following parameters can be used: D_*u*_ = 0.14, D_*v*_ = 0.06, f = 0.035, k = 0.065 and an example of the output is given in Figure 2. It is difficult to predict the pattern that will be produced for a given set of coefficients and some combinations will produce unstable outputs^64^. We find, at least for the system sizes that we have tested, that changing D_*v*_ while keeping the other coefficients constant will produce round ‘cells’, so long as D_*u*_ is at least twice as large as D_*v*_. After a stable pattern of spots is produced, a peak detection algorithm is then applied to the array (*v*) corresponding to the final time step, to identify the centres of each of the high contrast, well-defined cells that are formed.

**Figure 2.**
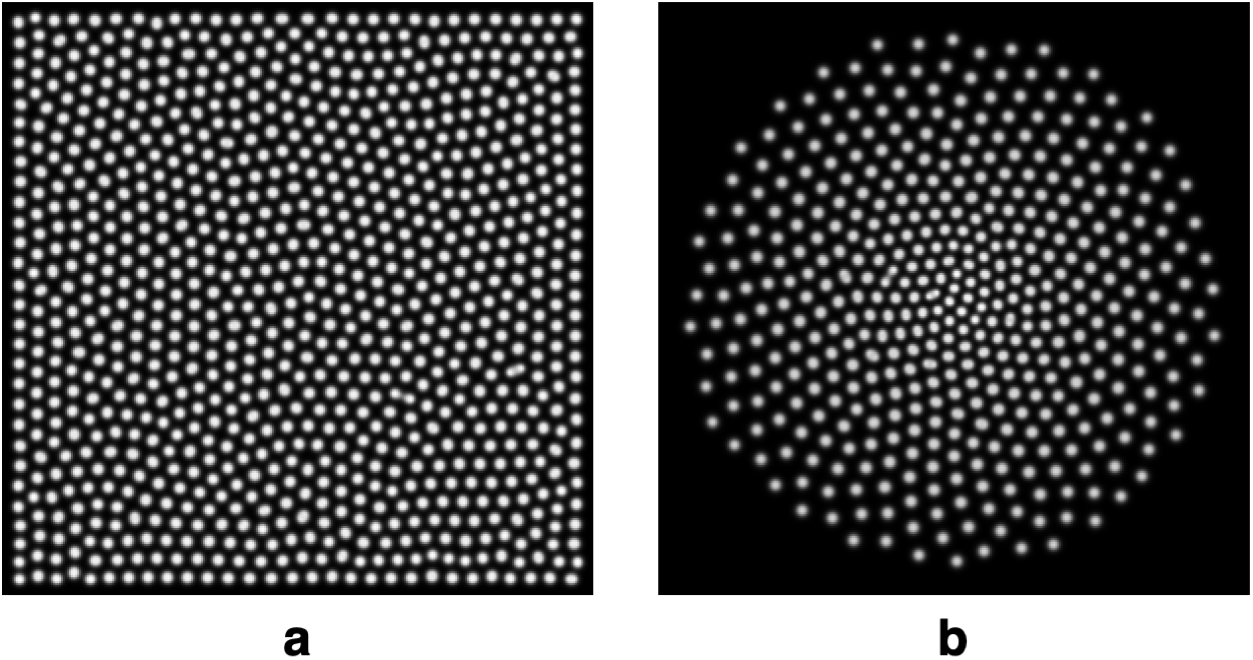
Self-organising mosaics generated using the Gray-Scott reaction-diffusion model. Panel **a** shows the output of a 300 x 300 element system after 20,000 time steps with coefficients: D_*u*_ = 0.14, D_*v*_ = 0.06, f = 0.035, k = 0.065. Panel **b** shows the output of a 300 x 300 element system after 60,000 time steps with coefficients: D_*u*_ = 0.14, D_*v*_ = 0.4 10^*−*3^r + 0.04, f = 0.035, k = 0.065, where *r* is the radial coordinate of the system, resulting in D_*v*_ = 0.04 in the centre and D_*v*_ = 0.12 at the furthest edge of the array. It should be noted that both models were seeded with the chemicals initially in the centre, and, in **b**, the simulation has been terminated before the chemicals have diffused to the edge of the system.

In the human eye, the spacing of the cone mosaic varies as a function of retinal eccentricity. The density of cones, *ρ*, can be approximated from histological measurements^65^ (raw data taken from^66^) and the spacing, *s*, calculated as^67^,

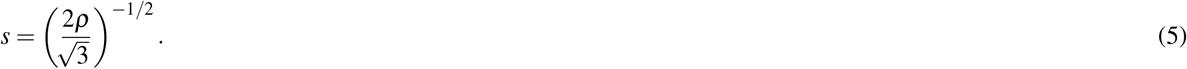

In principle, the spacing between the cells in the reaction-diffusion model can be tuned through selection of the appropriate diffusion coefficients, namely D_*u*_ and D_*v*_. Furthermore, defining a radial variation in these coefficients can produce a foveated mosaic of cells (see Figure 2**a**). However, care must be taken when selecting the diffusion coefficients, since the outcome (and its stability) will also depend on the coefficients *f* and *k*. The size of the system must also be taken into account to ensure that the sampling is sufficiently high for the spacing desired. For those reasons, it is more efficient to alter the cone spacing by simply re-scaling a well-sampled and stable cell mosaic. This is done within ERICA by generating a retinal mosaic (see Figure 2**a**) over an extended bounding box, and scaling the resulting cone positions such that the average cone spacing (determined by nearest neighbour analysis) matches the histological data.

#### Cone reflectance profiles

Photoreceptors act as optical waveguides^68^ and the number of guided modes (e.g. linearly polarised, LP, modes) depends on the physical properties of the cell. We consider each cone to have an intensity profile that can be approximated by a Gaussian function. Small cone cells close to the centre of the fovea (approximately <3°) act as single mode waveguides, and so this Gaussian model is reasonable. At larger eccentricities, cone cells exhibit biomodal (beyond 7°) or multimodal (beyond 10°) profiles and so the Gaussian approximation will be a less accurate representation as the support of higher-order modes results in more complex intensity profiles^69^. If simulation of such profiles is required ERICA could be modified to include higher-order modes in addition to the fundamental (Gaussian) mode, but clearly the modal composition should be validated against human-derived images. Furthermore, a full model of the light propagation through cone cells, including interference effects, that predicts the reflectance profiles seen in real AOSLO images has recently been developed by Meadway and Sincich^56^. More complex models such as these could be incorporated into the ERICA framework.

#### Mosaic ground truth data

An individual retinal mosaic is specified as an array of size N x M, where N is the number of cones in the mosaic and M is the number of parameters defining an individual cone’s elliptical Gaussian reflectance profile. As described above, cone locations extracted from the reaction diffusion model are scaled so that the spacing of the cells matches histological data. The sizes of the cells are specified as the diameter of the first (Gaussian) mode of the cell, which is calculated via the Marcuse equation with the cell’s radius determined from histological data^70,71^. ERICA specifies the following parameters for each cone in the mosaic: 1) the vertical coordinate, *µ*_*y*_, 2) the horizontal coordinate, *µ*_*x*_, 3) the vertical width, *σ*_*y*_, 4) the horizontal width, *σ*_*x*_, 5) the orientation, *θ*, and 6) the reflectance, *A*. This parameter array represents the ground truth for the make-up of the synthetic retina. The parameters can be defined by the user; aside from position, we use values randomly selected from a normal distribution based on histograms computed for real images. For cone reflectance, based on observations of real images, we additionally impose a low spatial frequency variation, and this approach has been previously described by others^53^. It is also be possible to specify temporal variations in the light reflected back from individual cones, such as might be seen on long time scales due to bleaching or disc shedding or on short time scales due to stimulus-evoked intrinsic changes in intensity^72^. In this case, the reflectance is taken to be that at the time point when the simulated scanned beam reaches the cone.

Using this approach it is possible to define a retinal mosaic analytically as a sum of Gaussians for *N* cones,

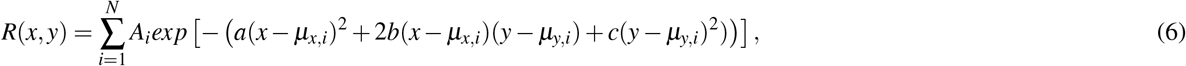

where

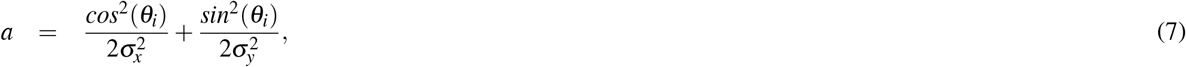

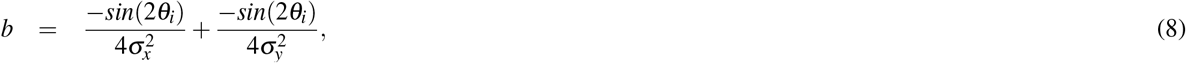

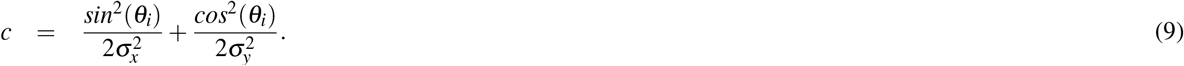

An analytic representation of the retina allows the AOSLO data capture, combining the scan pattern and the motion of the eye, to be replicated without introducing sampling errors. In the case where we wish to also replicate the effects of optical aberrations on the captured data it becomes necessary to perform additional computations, as described in a later section.

### Eye movement simulation

To introduce intra- and inter-frame motion, ERICA includes a biologically plausible model of the eye movements made during fixation. Fixational eye movements are typically categorized as either tremor, drift, or microsaccades^73,74^, each of which is incorporated into our model. The parameters defining each of these types of movement have default values, but these can be adjusted. Microsaccades are modeled as a ballistic motion with the peak velocity proportional to the amplitude of the displacement in log space. The amplitude of each microsaccade is randomly selected from a normal distribution (default: mean of 30 arcmin and a standard deviation of 3 arcmin^75^). In the model, the occurrences of microsaccades are pre-determined by randomly selecting time intervals from a normal distribution (default: mean of 1.5 seconds and a standard deviation of 0.25 seconds). Between the microsaccades, eye movement data are generated by combining tremor with drift. Tremor constitutes a high temporal frequency motion and is simulated using a random motion with a temporal bandpass filter^27^, which we model with a Gaussian profile (default: a central frequency of 65 Hz and a standard deviation of 7.5 Hz). The amplitude of tremor is typically very small^27^ and we model this by selecting it from a normal distribution with a low mean (default: a mean of 1 arcmin and a standard deviation of 0.1 arcmin). Drift is a low temporal frequency motion modeled by a 1/f^2^ spectrum, over which tremor is superimposed^76^. The amplitude for each period of drift is randomly selected from a normal distribution (default: a mean of 6.5 arcmin and a standard deviation of 0.7 arcmin). Human participants typically maintain fixation around a central point, however there has been debate over whether microsaccades or drifts are responsible for corrective motions to bring fixation back on target^77^. The directions of microsaccades in the ERICA model are constrained to ensure that the simulated fixational eye movement maintains a locus about the centre of the synthetic retinal mosaic, simply so that blank spaces from the edge of the mosaic are not introduced into the synthetic frames. An example eye movement trace is shown in Figure 3. The parameters outlined above can be adjusted by the user to generate eye movements with particular characteristics and, for example, to test how well those characteristics are recovered by a motion extraction algorithm.

**Figure 3.**
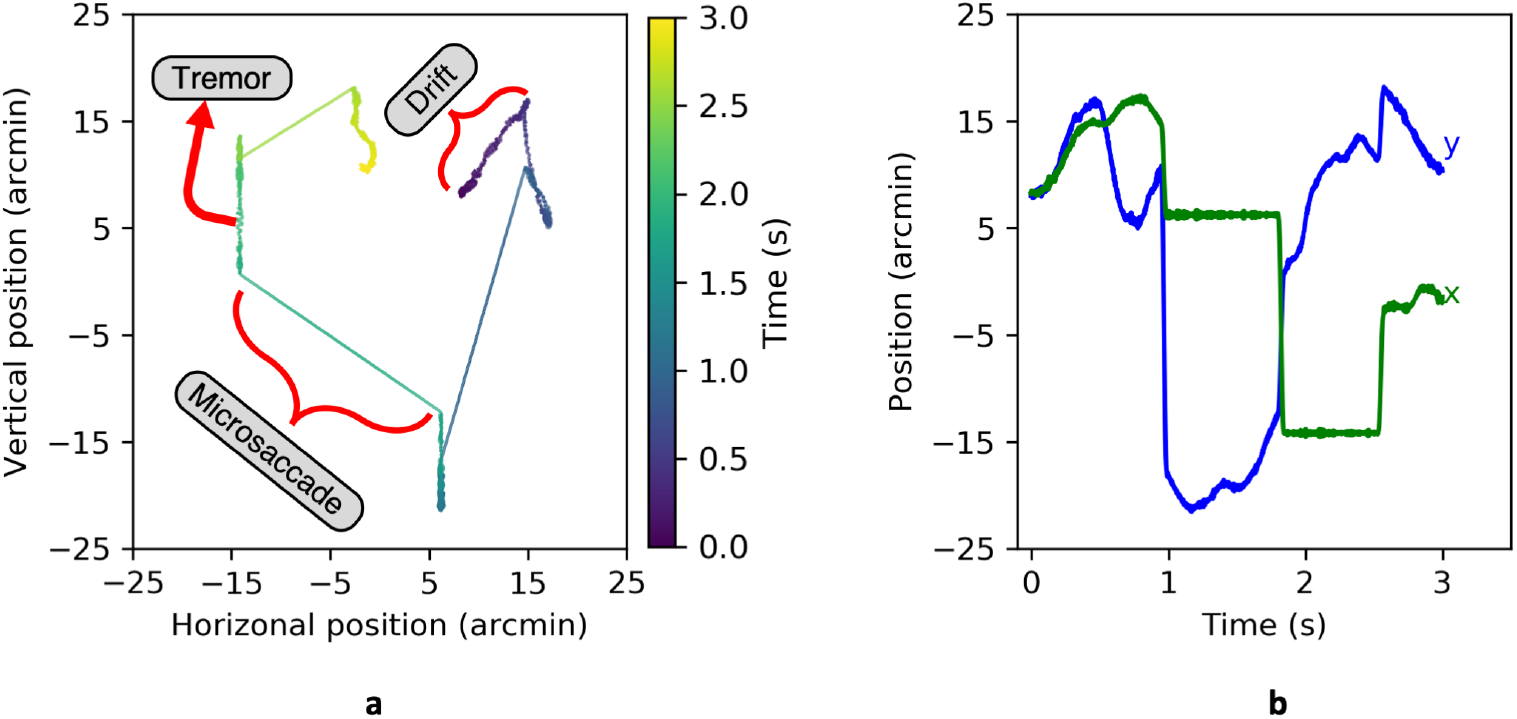
An example three-second eye movement trace, showing three microsaccades interspersed with periods of drift and tremor. Panel **a** shows the change in horizontal and vertical position over time, indicated by the colour gradient from blue (t_0*s*_) to yellow (t_30*s*_). Panel **b** shows the horizontal (x; green) and vertical (y; blue) components separately

### Image generation

The PSF of a double-pass confocal microscope, such as the AOSLO, is defined as the autocorrelation of the single pass PSF^78–80^. However, the retina is not a uniform diffuse reflector. To replicate the process of data capture from the retinal mosaic a series of computations must be performed as described in Figure 4. Briefly: 1) the input PSF should be multiplied by the synthetic retinal mosaic, giving the reflectance profile of the illuminated patch of retina; 2) the illuminated patch should be convolved with the output PSF to replicate reimaging it on to the pinhole; and 3) the image formed at the pinhole should be multiplied by the transmission function defining the pinhole. The sum of this final image is the signal that would be received at the detector at a particular moment in time. It is possible to perform these computations by adding terms and operations to Equation 6, such as by using the extended Nijboer-Zernike approach to approximate the analytic PSF^81,82^. However, this adds complexity and that method also assumes some wavefront symmetry^81^. It is therefore simpler to use a numerical approach and use the discrete Fourier transform to render images with residual aberration (which can vary from frame-to-frame), taking care to oversample appropriately to avoid sampling artefacts. Finer sampling resolutions produce images closer to an analytic solution but with added computational expense. To test this, a set of retinal mosaics was generated with a range of sampling resolutions and this set was used to generate synthethic frames with a realistic residual wavefront error (based on a publicly available data set^61^). A sampling resolution at least two times higher than the desired resolution of the resulting frame should be used and the test showed that a factor increase in sampling resolution of *≥*4 produces almost identical outputs, defined by the sum squared difference from the a high (10*×*) sampling resolution image.

**Figure 4.**
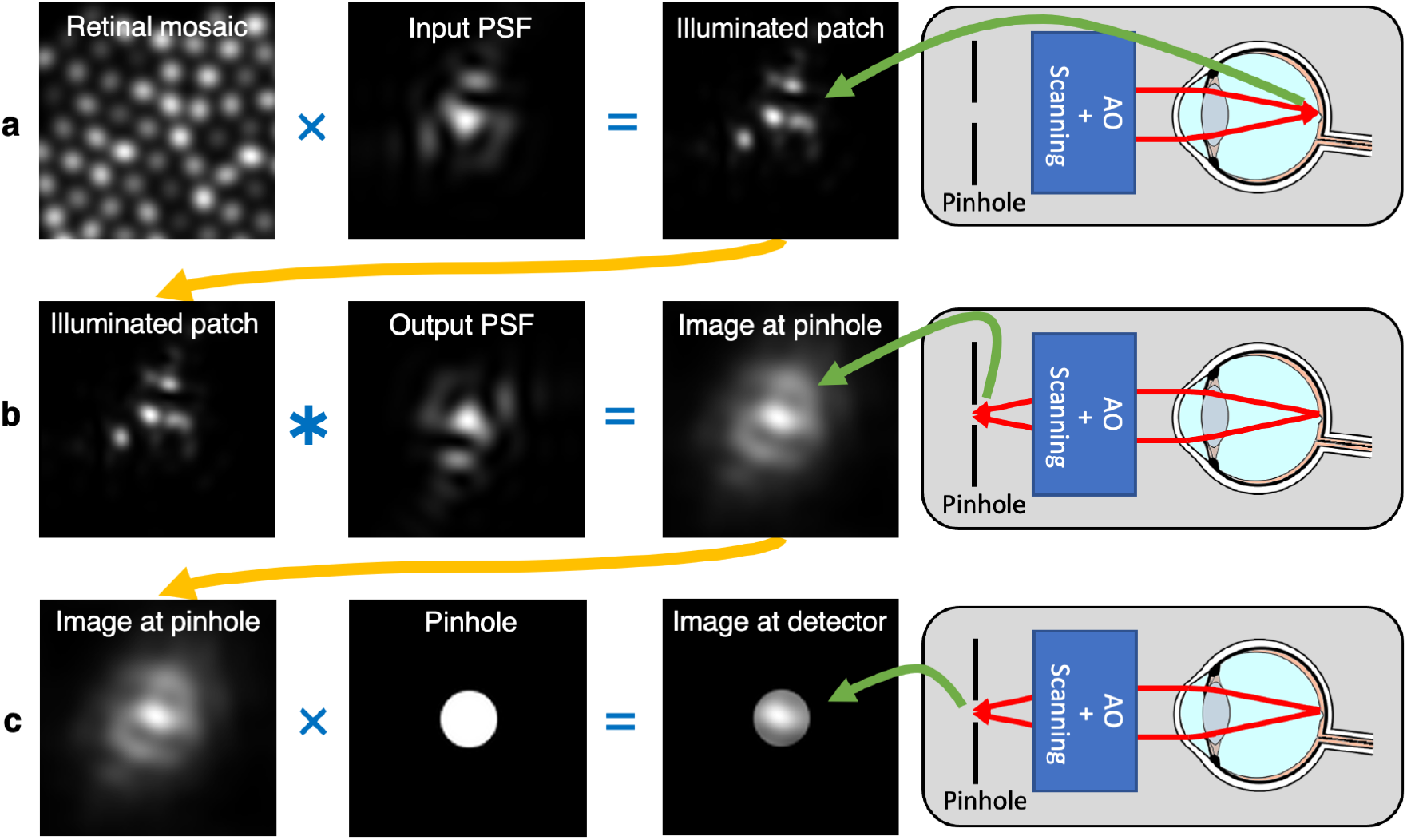
The steps to replicate data capture through a scanned confocal retinal imaging system, such as an AOSLO, which are shown schematically on the right. Step **a** shows the illumination of a patch of retina by a point source directed through the system and the eye’s optics (Input PSF), which may be degraded by the eyes optics. This step is represented by the multiplication of the reflectance profile of the retinal patch and the input PSF. Step **b** shows the process of re-imaging the illuminated patch of the retina through the system and the eye’s optics (Output PSF) to the plane where the pinhole is located. This step is computed as a convolution of the illuminated patch of retina with the output PSF, noting that the wavefront is inverted in the outward direction (this operation is equivalent to the cross-correlation of the illuminated patch with the input PSF). Step **c** shows the transmission of the final image through the pinhole to give the image that would be seen by the detector. This step is computed as the multiplication of the image at the pinhole by the transmission function of the pinhole, which we consider to be 1 inside and 0 outside a uniform disc. The final step is summation of the image formed behind the pinhole, to replicate integration by the detector (neglecting noise), giving the signal at the current position in the scan, *I(x,y)*. Steps **a-c** are repeated for every sample location as the beam is raster scanned over a region of the retina. Note that while this model allows for a different input and output PSF, in practice we assume that the input and output PSFs are the same (the PSF of the eye, which includes diffraction effects).

Following the steps in Figure 4, ERICA generates a data stream similar to that which would be obtained from an AOSLO (or any scanned confocal system, such as a standard scanning laser ophthalmoscope). A raw image frame is then generated from the data stream using knowledge of the positions on the scanning mirrors at each time point. To generate a motion free reference image, a data stream is generated by replicating the spatio-temporal sampling of the raster scanning system - a sinuoidal horizontal fast scan and an approximately linear vertical slow scan. To generate a realistic sampling pattern during fixation, eye movement would be added to the raster scan pattern, as described above. The scan pattern used in the simulation can be defined by the user and we use a model of the scanning mirrors in our AOSLO^83^. This data stream is then typically processed using the same desinusoiding algorithm that is applied to real, human-derived intensity data. All images presented in this paper have been desinusoided for clarity. Noise can be added to images after following the steps in Figure 4 and we assume that this is an additive process. By default, noise is generated by selecting values from a normal distribution, the mean and standard deviation of which are matched to the noise characteristics of our AOSLO. This array of noise samples must be either added before desinusoiding, or must must undergo the same desinusioding procedure before being added to the image.

### Comparison to human cone mosaics

To verify that the packing geometry of the synthetic mosaics is representative of the cone mosaic of typical human participants, we made comparisons at a set of matched retinal eccentricities. Images were collected from seven human participants at three eccentricities (1.5°, 3.0° and 6.0°) as part of another study, which received approval from the University of Oxford Research Ethics Committee (reference: R57315/RE001). The study was conducted in accordance with the Declaration of Helsinki and informed consent was obtained from all participants. We did not attempt imaging in the centre of the fovea as without dilation the cone cells would not be well-resolved and the estimates of cone packing would likely be unreliable. Cone positions were manually labeled in both the human and synthetic images by five experienced graders (one of whom was the first author) and metric data were averaged across graders. Manual grading of the synthetic images, rather than using the ground truth data, ensured that apparent changes in the packing geometry caused by weakly reflecting cones, which could be missed by a human grader, were captured for a fair comparison to the human images. Presentations of synthetic and human images at different eccentricities were randomly interleaved and the grader was blind to the type of image displayed on any given trial.

### Extensions

The current implementation of ERICA simulates the cone mosaic as visualised through a scanning confocal imaging system. This model is limited to a 2D representation of the retina and residual wavefront aberrations are added based on a 2D model of the PSF. Modelling the effects of changing the axial resolution would be possible with an extension of the simulation to perform image generation based on the 3D PSF and with a cellular-resolution model of the retina in depth. A model of the light propagation through the cone cells, such as that developed by Meadway and Sincich could also be incorporated into the ERICA framework.

A simpler approach to generating images of other retinal structures, either in depth or within the same retinal layer, is to estimate the intensity profile of those structures. Examples are given in Figure 5. Figure 5**a** shows that rod cells could be added into the images. This is possible by modification of the reaction-diffusion model to self-organise smaller cells in amongst a mosaic of larger cells. This will be presented in more detail in a future publication. Another structure that is likely to be of interest is the retinal vasculature, particularly as it causes shadows that may affect image registration. A basic approximation, as shown in Figure 5, would be to introduce a darkened region, either during or after image generation. Non-confocal imaging geometries detect multiply-scattered light returning from structures other than the photoreceptors. Accurately replicating this process would require a ray-tracing model, which is not currently included within in ERICA. However, it is possible to generate approximate non-confocal synthetic images using an estimate of the intensity profile of the structures observed in these images. As an example, split-detection imaging is a useful tool for detecting photoreceptor inner segments even when the reflectance in the confocal image is low. Methods for automatic detection of cells in these images and for montaging have been developed^41,57,58^. An example of a synthetic split-detection image is given in Figure 5 **c**, which has been generated by approximating the expected intensity profile as the differential of a Gaussian distribution. The example given shows differentiation in the horizontal direction, replicating a left-right split-detection geometry. Similarly, a model image of the retinal pigment epithelium (RPE), is theoretically possible, such as by using the reaction-diffusion model to generate cell locations and using an estimated dark-field RPE cell intensity profile. By updating the ground truth retina, ERICA allows emulation of the AOSLO image capture from such structures, including realistic eye motion, aberrations and noise.

**Figure 5.**
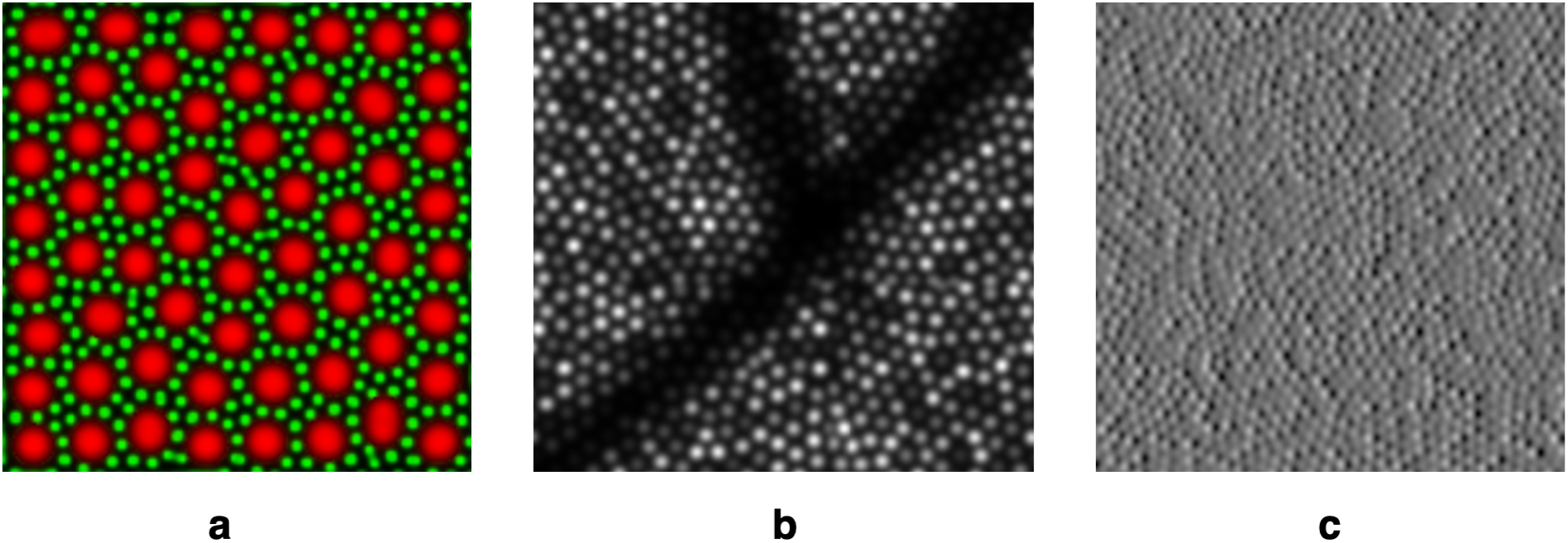
Examples to show possible extensions to ERICA. Panel **a** shows the output of a reaction-diffusion model that includes a rod mosaic (shown in green), self-organising itself between cone cells (shown in red). Panel **b** shows simulation of the shadow caused by a retinal blood vessel overlaid on a synthetic cone mosaic image. Panel **c** shows the approximation of a split-detection image, formed by reproducing the intensity profile of a photoreceptor inner segment as it typically appears in such images. This is assumed to be a differential image of a mosaic of Gaussian profiles.

## Results

Examples of synthetic images produced by ERICA are given in Figure 6 and an example movie showing temporal variation in eye motion and wavefront error, based on real aberration measurements^61^, is additionally given in the supplementary materials. The appearances of the synthetic and human images are convincingly similar. The lower panels in Figure 6 show the normalised power spectra of the images in the top panels. The spacing of the cells is matched within ERICA based on the retinal eccentricity and the axial length of the participant’s eye and so the modal spacing (the peak in the power spectrum at <50 c/deg, which corresponds to Yellott’s ring) is also matched. Therefore, the scale of the spatial frequency axis is matched between human and synthetic images, but these spectra also show similar profiles.

**Figure 6.**
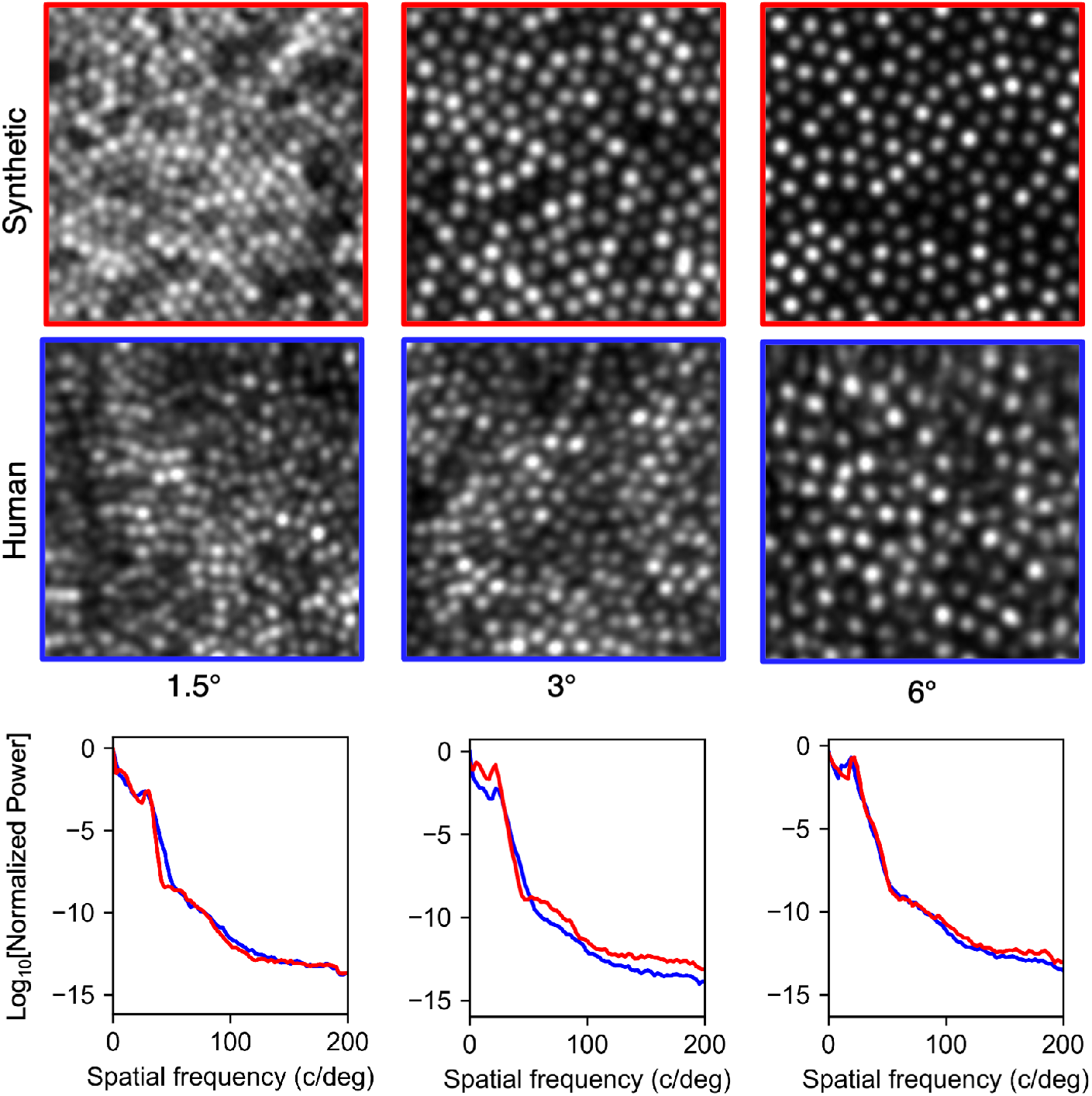
Example images from a human participant collected from our AOSLO^83^ and synthetic images generated for the same eccentricities - 1.5°, 3° and 6°. The synthetic images are generated with no noise, residual aberration or eye motion and so have better image quality than the human images. The pupil diameter for the simulation was set to 5 mm, which is approximately the same as the diameter of the human participant’s eye during image capture. The synthetic images were spatially scaled to account for the human participant’s axial length being different to that of the model. The synthetic images presented here have been intensity-scaled linearly and then clipped such that the number of pixels that are saturated is the same as in the human images (approximately 10% for the detector gain that was used during imaging). The lower panels show the power spectra of the synthetic images in red and the human images in blue. In each spectrum, the peak below 50c/deg corresponds to the modal spacing of the cones (Yellot’s ring).

We analysed the packing geometry of the synthetic and human cone mosaics using three summary statistics (see Figure 7): a) the number of nearest neighbours to a cone (the number of Voronoi regions associated with an individual cone), b) the first nearest neighbour distance, and c) the vertex angle between the nearest neighbour and that neighbour’s closest common neighbour). For each image and at each eccentricity, the three parameters were calculated for 100 randomly selected cones. The metric value for an individual image was defined as the standard deviation of the calculated parameter, normalised by the mean (or 60° in the case of the vertex angle measurement). This gives a measure of variance that would be zero for a perfectly regular hexagonal matrix. Data were then averaged across images (across participants) for each eccentricity. To help interpret the scale of these values, a cone mosaic was generated with a random packing arrangement (but the same number of cells) and without any constraint on the minimum spacing between cells. This provides an estimate of the metric value for a completely irregular mosaic. Figure 7 shows a comparison between the packing regularity of synthetic and human images. These results show that ERICA produces cone packing geometries close to those that we observe in the human retina through our AOSLO^83^. It is notable that agreement between the human images and the simulation is reduced at the largest eccentricity when considering the variation in nearest neighbour distance. We suggest this is likely due to the presence of rods in the human images that are absent in the simulation, which may have affected the manual labels^84^.

**Figure 7.**
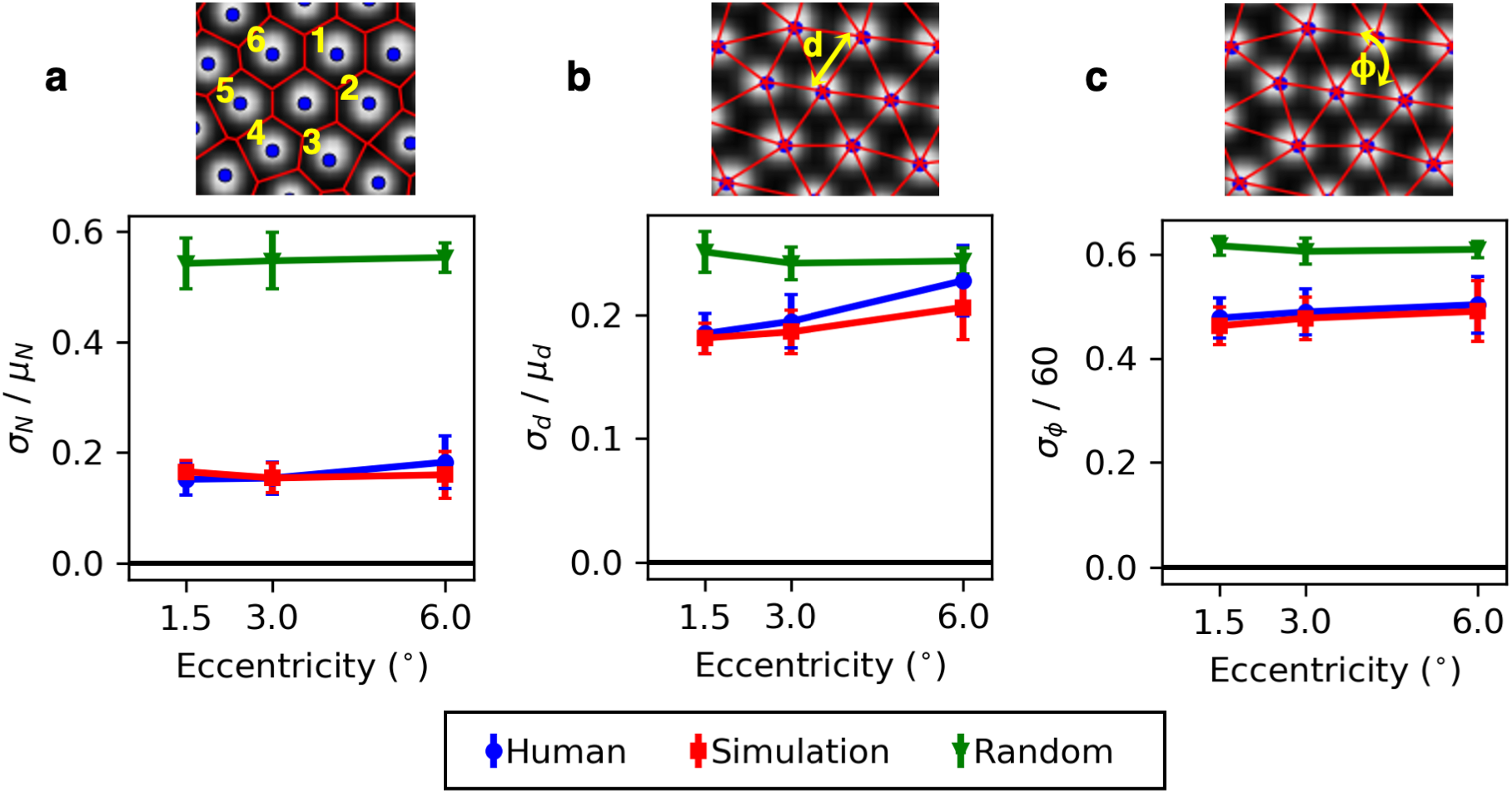
Three metrics of packing regularity (eccentricities the same as in Figure 6) are compared for human-derived images and for synthetic images generated either using reaction-diffusion modelling or by randomly placing cells (i.e completely irregular packing with the same number of cones). For each metric the standard deviation is used as a measure of variability, normalised by the mean. A value of zero is equivalent to a completely regular mosaic. Each data point represents the mean, and each error bar the standard deviation, calculated over 27 human or 12 synthetic images. Images were manually labeled by five experienced markers and the data plotted here represent the average across markers. The panels show the variability in **a** the number of nearest neighbours, **b** the nearest neighbour distance and **c** the vertex angle between nearest neighbours.

## Discussion

Adaptive optics retinal imaging is rapidly emerging as an important tool for studying the eye^4,5,85^. A significant bottleneck in the wider use of these systems is the need for robust, accurate and automated image processing and analysis methods to generate good quality images and extract reliable data. The use of a generative model for algorithm testing is not new, but its implementation in the AO retinal imaging field has so far been limited^52–55^. ERICA uses a self-organising approach that produces mosaics with similar packing geometries to human-derived images. An advantage to using reaction-diffusion modelling is that cells naturally arrange themselves without gaps. Self-organising approaches has been suggested previously as a model for retinal sampling^62,86^, using different methods. However, the reaction-diffusion approach does offer another advantage, as demonstrated in Figure 5**a**, since multiple interleaved mosaics can be generated that both self-avoid and avoid each other.

Previous approaches for generating synthetic images have performed image generation in a similar way to the method we have developed. These approaches model cone cells as Gaussian intensity profiles and convolve the retinal image with the PSF^53,54^. ERICA fully replicates the image generation process, as shown in Figure 4, and we include a step to separately specify the input and output PSFs to include, for example, differences in the input and output pupil diameters.

Beddgood and Metha developed a model to add motion distortions to simulated images using a different approach to ours^50^. They used a flood-illumination image as the ground truth and the rows of this were re-indexed based on an eye movement trace that had been captured from a human participant. By contrast, ERICA includes a model of biologically plausible eye motion allowing motion extraction accuracy to be characterised relative to the specific measures of interest, such as the amplitude or velocity of microsaccades or drifts. These movements are introduced during a replication of the non-linear scan pattern and this can additionally be performed analytically to avoid sampling artefacts. De-sinusoiding is a necessary step for processing images from an AOSLO (and from a scanning laser ophthalmoscope without AO correction). Dependent on the voltage ramp of the galvo, it is also possible that the slow scanner is not perfectly linear. By including a model for the scan pattern we can test the effect that non-linearities have on motion extraction, in terms of both the remapping of spatial information and the temporal sampling of eye motion, which ideally would be linear.

In this paper we have presented a complete end-to-end simulation of AOSLO data capture that allows generation of large data sets of retinal images with ground truth data. The properties of these synthetic images, such as residual wavefront error, noise and motion distortion, as well as the content (e.g. by varying retinal eccentricity or changing cone reflectances) can be specified. ERICA has been designed in a modular framework so that any part of the simulation could be extended or modified, and any parameters adjusted, in order to replicate a user’s own system. Finally, we openly share ERICA with the community and make a number of synthetic data sets available. We hope that this will be a useful contribution to the field to further the design, development and testing of reliable automated image processing and analysis methods for adaptive optics retinal imaging.

## Acknowledgements

The authors would like to thank Sarah Regan and Mital Shah for collecting the human images and the five expert raters for producing manual labels for them. The authors would like to acknowledge funding from Fight For Sight (1467/8), the University of Oxford Wellcome Trust Institutional Strategic Support Fund (105605/Z/14/Z), the University of Oxford Medical Research Fund (MRF/LSV2015/2161), the EPA Cephalosporin Fund (CF 277), the John Fell Oxford University Press (OUP) Research Fund (103/786 and 151/139). Laura Young is supported by a UKRI Future Leaders Fellowship (MR/T042192/1) and by the Reece Foundation.

## Author contributions statement

Laura Young designed and wrote the simulation, performed the analyses and wrote the main manuscript text. Both authors contributed to the conceptualisation of the approach and to writing and editing the manuscript.

## Additional information

ERICA is available to the community via GitHub (https://github.com/LauraKateYoung/ERICA). Example data sets are also available for use (https://data.ncl.ac.uk/authors/Laura_*Y*_*oung/*9553463).

