## Supplemental text for "ERICA: Emulated Retinal Image CApture - A tool for testing, training and validating retinal image processing methods"

### Supplementary materials

Supplementary Video 1 shows a sequence of 60 synthetic frames generated using ERICA that constitute a 2 second movie. Encoded in the frames are variations in image quality due to residual aberrations not corrected by the optical system, which are modeled using a a dataset of wavefront measurements captured from real eyes[1]. The synthetic frames have been generated with modeled movements of the eye that include drift and tremor but no microsaccades.
